# Generation of three induced pluripotent stem cell lines from an immune checkpoint inhibitor-induced myocarditis patient and controls

**DOI:** 10.64898/2026.06.17.730743

**Authors:** Yin Sun, Maria Rosaria Vitale, Anna P. Hnatiuk, Noah S. Wagner, Shuai Sun, Xiaochun Yang, Lu Liu, Shaheen Khatua, Hiranya Amritavalli Sundar, Harrison Chou, Yuhsin Vivian Huang, Sarah Waliany, Yan Zhuge, Ronald Witteles, Mark Mercola, Joseph C. Wu, Han Zhu

## Abstract

Immune checkpoint inhibitor–associated myocarditis (ICIM) is an uncommon but potentially fatal inflammatory heart disease triggered by cancer immunotherapy, with up to 40% mortality. The underlying mechanisms are still elusive, partly due to the lack of appropriate human disease models. Here, we report the generation of three induced pluripotent stem cell (iPSC) lines derived from an ICIM patient, an ICI-treated patient without myocarditis, and a healthy donor. These lines exhibit typical pluripotent stem cell morphology, express pluripotency markers, maintain normal karyotypes, and differentiate into derivatives of the three germ layers, providing a valuable platform for mechanistic studies and therapeutic discovery.

## Resource Table

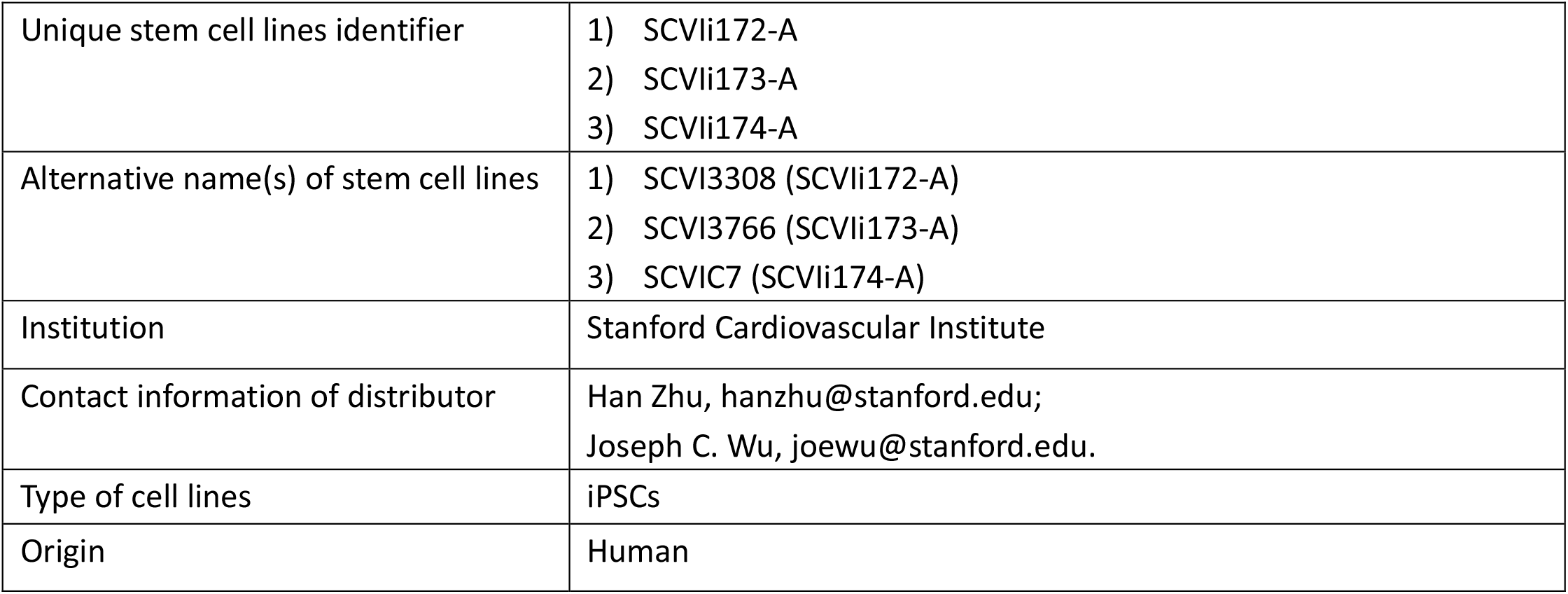
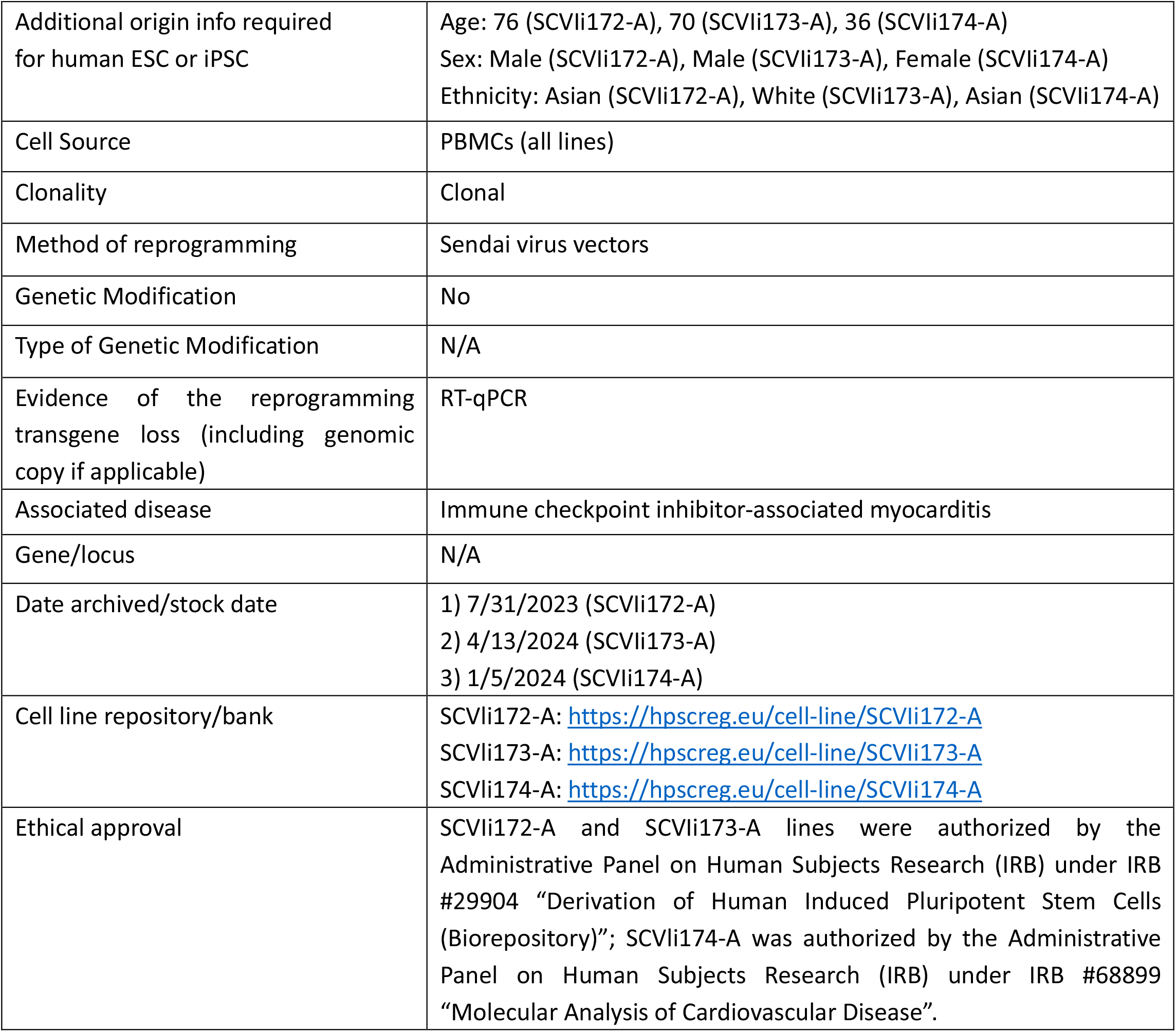

## Resource utility

The three induced pluripotent stem cell (iPSC) lines in this study were generated from a patient with ICIM, an ICI-treated patient without myocarditis, and a healthy donor. The iPSC lines provide a resource to model ICIM for mechanistic studies and drug discovery. The inclusion of both an ICI-exposed control without myocarditis and a healthy donor provides important controls to test disease-specific susceptibility. Although ICIM is a treatment-associated complication, individual susceptibility may be influenced by patient-specific immune features, such as HLA-dependent antigen presentation and T cell recognition. iPSCs retain donor-specific genomic information and can be differentiated into cardiomyocytes. Importantly, these lines enable matched or HLA-compatible co-culture systems with immune cells, providing a platform to study T cell–mediated cardiotoxicity in ICIM. Therefore, patient-derived iPSC lines provide a valuable platform to model disease-relevant cellular phenotypes.

## Resource Details

Immune checkpoint inhibitors (ICIs) have revolutionized cancer therapy but can induce immune checkpoint inhibitor–associated myocarditis (ICIM), a rare yet life-threatening complication with a mortality rate of up to 40% (Zhu et al., 2022; Huang et al., 2025; Johnson et al., 2016). Despite its clinical significance, the cellular and molecular mechanisms underlying ICIM remain largely elusive, due to the lack of appropriate human disease models that faithfully recapitulate patient-specific immune and cardiac interactions (Waliany et al., 2021). Induced pluripotent stem cells (iPSCs) provide a valuable platform for modeling human diseases, as they retain patient-specific characteristics and can be differentiated into relevant cell types, including cardiomyocytes and immune cells (Sun et al., 2026; Wu et al., 2025). Therefore, the establishment of disease-specific iPSC lines represents a valuable approach to investigate the pathogenesis of ICIM and to enable the development of targeted therapeutic strategies.

In this study, we established three iPSC lines (SCVIi172-A, SCVIi173-A, and SCVIi174-A) from peripheral blood mononuclear cells (PBMCs) using Sendai virus-mediated transduction of Yamanaka factors. SCVIi172-A was derived from a 76-year-old Asian male, who received ICI treatment (Durvalumab) and diagnosed with ICIM, SCVIi173-A from a 70-year-old white male who received ICI treatment (Durvalumab) without developing myocarditis, and SCVIi174-A from a 36-year-old Asian female healthy donor. All three lines exhibited typical iPSC morphology (Fig. 1A; scale bar: 130 μm). The undifferentiated iPSC state was detected using pluripotency markers (SOX2, NANOG, and OCT3/4) by immunofluorescence (IF) staining (Fig. 1B; scale bar: 100 μm). Moreover, the expression of *SOX2* and *NANOG* at the mRNA level was validated by quantitative reverse transcription polymerase chain reaction (RT-qPCR) (Fig. 1C), showing robust expression in all iPSC lines but not in fibroblasts (negative control). Quality control analyses demonstrated that all lines were free of mycoplasma contamination and Sendai virus (Fig. 1D and E, respectively). Short tandem repeat (STR) analysis confirmed that the genetic profiles of the iPSC lines matched those of the original donor PBMCs (data submitted to Journal archive). Furthermore, all three lines successfully differentiated into derivatives of the three germ layers—ectoderm, endoderm, and mesoderm—as confirmed by immunostaining for OTX2/PAX6, SOX17/FOXA2, and BRACHYURY/TBX6, respectively (Fig. 1F; scale bar: 100 μm). KaryoStat assay suggested that all three lines maintained a normal karyotype (Fig. 1G).

**Fig. 1.**
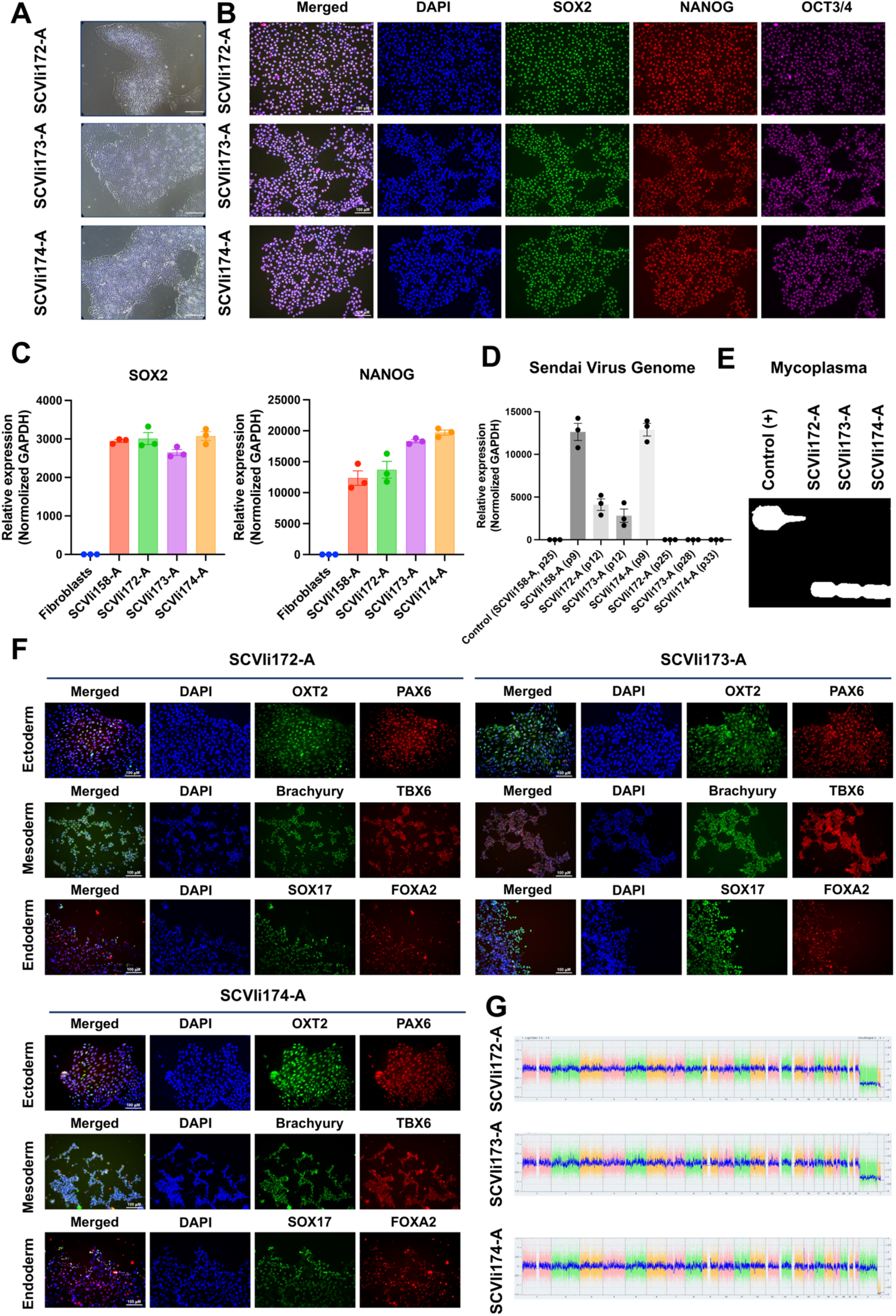
Characterization of three iPSC lines: SCVli172-A (68899), SCVli173-A (68899), and SCVli174-A.

## Materials and Methods

### 1. Reprogramming

PBMCs were isolated from whole blood of a patient with ICIM, an ICI–treated patient without myocarditis, and a healthy donor using Percoll density gradient medium (GE Healthcare #17089109), followed by washing with DPBS (Thermo Fisher Scientific #14190144). PBMCs were cultured in StemPro®-34 SFM medium (Thermo Fisher Scientific #10639011) supplemented with 20 ng/mL IL-6 (#PHC0063), 20 ng/mL IL-3 (PeproTech #200-03), 20 ng/mL EPO (Thermo Fisher Scientific #PHC9631), 100 ng/mL FLT3 ligand (#PHC9414), and 100 ng/mL SCF (PeproTech #300-07). Reprogramming was performed using the CytoTune™-iPSC 2.0 Sendai Reprogramming Kit (Thermo Fisher Scientific #A16517) according to the manufacturer’s instructions, with 5 × 10^5^ PBMCs per well used for Sendai virus-mediated transduction. During transduction, cells were plated onto Matrigel-coated plates. Seven days post-transduction, cells were cultured in StemMACS™ iPS-Brew XF medium (Miltenyi Biotec #130-104-368) until colonies emerged. Colonies typically appeared 10-15 days post-transduction and were manually picked, expanded, and cryopreserved for further experiments.

### 2. Cell culture

All cultures were maintained at 37°C and 5% CO_2_ in a humidified incubator. Cells were cultured on Matrigel-coated plates (1:500, Corning #356231) in mTeSR™ Plus medium (STEMCELL Technologies 05826) containing mTeSR™ Plus 5X Supplement (STEMCELL Technologies 05827). For maintenance, the medium was changed the day after passaging and every other day. For passaging, cells were dissociated using 0.5 mM EDTA (Invitrogen #15575-038) and reseeded in medium containing 10 *μ*M ROCK inhibitor (Y-27632, Selleck Chemicals #S1049) for the first 24 hours. All assays were performed using iPSCs between passages 25 and 30, unless otherwise specified.

### 3. RNA extraction and RT-qPCR

Expression of pluripotency markers (*SOX2* and *NANOG*) and Sendai virus clearance were assessed by RT-qPCR. Sendai virus clearance was assessed at both early (P9-P12) and later passages (≥ P25) to confirm the complete clearance of Sendai viral vectors. Total RNA was extracted from iPSC lines using TRIzol reagent (Thermo Fisher Scientific) according to the Direct-zol RNA Microprep Kit protocol (Zymo Research #R2062). Fibroblasts were used as a negative control. A previously characterized human iPSC line (SCVIi158-A), generated by the Dr. Joseph C. Wu laboratory (Sun et al., 2026), was used as a reference control for pluripotency marker expression and as a positive control for Sendai virus detection in RT-qPCR assays. cDNA was synthesized using the iScript cDNA Synthesis Kit (Bio-Rad #1708891), and quantitative PCR was performed using predesigned TaqMan probes (Table 2).

**Table 1.** Characterization and validation.

| **Classification** | **Test** | **Result** | **Data** |
| --- | --- | --- | --- |
| **Morphology** | Photography Bright field | Visual record of the line: normal | Figure. 1 panel A |
| **Characterization of undifferentiated state** | Quantitative analysis: Immunofluorescence staining | Positive expression of undifferentiated iPSC state markers: OCT3/4, NANOG, and SOX2 in the three iPSC lines | Figure. 1 panel B |
|  | Qualitative analysis: RT-qPCR | Positive expression of undifferentiated iPSC state markers *NANOG* and *SOX2* in the three iPSC lines and absent in fibroblasts | Figure. 1 panel C |
| **Genomic stability** | Whole genome array (KaryoStat™ Assay) resolution | Normal karyotype: 46, XY for line SCVli172-A and SCVli173-A, and 46, XX for line SCVli174-A | Figure. 1 panel G |
| **mtDNA analysis (IF APPLICABLE)** | Sanger sequencing, Deep sequencing, analysis software (Mitopore www.mitopore.de, mtDNA-Server 2, etc) | N/A | N/A |
| **Cell line identity verification** | STR analysis | 16 loci tested match well | Available with the authors |
|  | Microsatellite PCR (mPCR) OR SNP array | N/A | N/A |
| **Mutation analysis (IF APPLICABLE)** | Sequencing | N/A | N/A |
|  | Southern Blot OR WGS | N/A | N/A |
| **Microbiology and virology** | Mycoplasma | Mycoplasma testing luminescence: Negative | Figure. 1 panel E |
| **Characterization of the differentiated state** | Directed tri-lineage differentiation | Positive IF staining of three germ layer markers | Figure. 1 panel F |
| **List of recommended differentiated state germ layer markers** | Expression of these markers has to be demonstrated at mRNA (RT PCR) or protein (IF) levels, at least 2 markers need to be shown per germ layer | Ectoderm: PAX6, OTX2<br>Endoderm: SOX17, FOXA2<br>Mesoderm: Brachyury, TBX6 | Figure. 1 panel F |
| **Donor screening (OPTIONAL)** | HIV 1 + 2 Hepatitis B, Hepatitis C | N/A | N/A |
| **Genotype additional info (OPTIONAL)** | Blood group genotyping | N/A | N/A |
|  | HLA tissue typing | N/A | N/A |

**Table 2.** Reagents details.

| **Antibodies used for immunocytochemistry/flow-cytometry** |  |  |  |  |
| --- | --- | --- | --- | --- |
|  | **Antibody** | **Dilution** | **Company Cat #** | **RRID** |
| Undifferentiated iPSC state marker | Rabbit Anti-Nanog | 1:200 | Proteintech<br>Cat #142951-1-AP | RRID:<br>AB_1607719 |
| Undifferentiated iPSC state marker | Mouse IgG2bk Anti-Oct-3/4 | 1:200 | Santa Cruz Biotechnology<br>Cat #sc-5279 | RRID:<br>AB_628051 |
| Undifferentiated iPSC state marker | Mouse IgG1k Anti-Sox2 | 1:200 | Santa Cruz Biotechnology<br>Cat #sc-365823 | RRID:<br>AB_10842165 |
| Ectoderm marker | Rabbit Anti-Pax6 | 1:200 | Thermo Fisher Scientific<br>Cat #42-6600 | RRID:<br>AB_2533534 |
| Ectoderm marker | Goat Anti-Otx2 | 1:200 | R&D Systems<br>Cat #963273 | RRID:<br>AB_2157172 |
| Mesoderm marker | Goat Anti-Brachyury | 1:200 | R&D Systems<br>Cat #963427 | RRID:<br>AB_2200235 |
| Mesoderm marker | Rabbit Anti-Tbx6 | 1:200 | Thermo Fisher Scientific<br>Cat #PA5-35102 | RRID:<br>AB_2552412 |
| Endoderm marker | Goat Anti-Sox17 | 1:200 | R&D Systems<br>Cat #963121 | RRID:<br>AB_355060 |
| Endoderm marker | Rabbit Anti-Foxa2 | 1:200 | Thermo Fisher Scientific<br>Cat #701698 | RRID:<br>AB_2576439 |
| Secondary antibody | Alexa Fluor 488<br>Goat Anti-Mouse IgG1 | 1:1000 | Thermo Fisher Scientific<br>Cat #A-21121 | RRID:<br>AB_2535764 |
| Secondary antibody | Alexa Fluor 488<br>Donkey Anti-Goat IgG (H + L) | 1:1000 | Thermo Fisher Scientific<br>Cat #A-11055 | RRID:<br>AB_2534102 |
| Secondary antibody | Alexa Fluor 555<br>Goat Anti-Rabbit IgG (H + L) | 1:500 | Thermo Fisher Scientific<br>Cat #A-21428 | RRID:<br>AB_141784 |
| Secondary antibody | Alexa Fluor 647<br>Goat Anti-Mouse IgG2b | 1:250 | Thermo Fisher Scientific<br>Cat #A-21242 | RRID:<br>AB_2535811 |
| **Primers** |  |  |  |  |

|  | Target | Size of band | Forward/Reverse primer (5'-3') |
| --- | --- | --- | --- |
| Sendai virus plasmids (qPCR) | Sendai virus genome | 181 | Mr04269880_mr |
| House-keeping gene (qPCR) | *GAPDH* | 471 | Hs02786624_g1 |
| Undifferentiated iPSC state marker (qPCR) | *SOX2* | 258 | Hs04234836_s1 |
| Undifferentiated iPSC state marker (qPCR) | *NANOG* | 327 | Hs02387400_g1 |

### 4. Immunofluorescent staining

Cell lines were fixed with 4% paraformaldehyde (Fisher Scientific #50980487) for 15 minutes, then permeabilized with 0.1% PBST (PBS containing 0.1% Triton X-100, Sigma-Aldrich #9036-19-5) for 10 minutes. Next, Cells were then blocked with 5% goat serum (#G9023) or 5% donkey serum (#D9663) in PBST for 30 min, depending on the host species of the secondary antibody. Primary antibodies (Table 2) were incubated overnight at 4 °C, followed by incubation with corresponding secondary antibodies for 30 min at room temperature. Nuclei were counterstained with DAPI (Thermo Fisher #P36931), and images were acquired using an ECHO Revolve microscope.

### 5. Trilineage differentiation

The differentiation potential of iPSC lines into the three germ layers was assessed. For endoderm differentiation, cells were induced using the STEMdiff™ Definitive Endoderm Differentiation Kit (STEMCELL Technologies #05110) according to the manufacturer’s instructions. For mesoderm differentiation, cells were treated with RPMI medium supplemented with B27 minus insulin (Gibco #11875-085 and #A18956-01) and 6 *μ*M CHIR99021 (Selleck Chemicals #S2924) for 48 h. For ectoderm differentiation, cells were induced using the Human Pluripotent Stem Cell Functional Identification Kit (R&D Systems #SC027B). Differentiation into all three germ layers was confirmed by immunofluorescent staining of lineage-specific markers.

### 6. Short tandem repeat analysis

Genomic DNA was extracted from iPSCs and their originating PBMCs using the DNeasy Blood & Tissue Kit (Qiagen #69504). STR profiling was performed at the Stanford Protein and Nucleic Acid (PAN) Facility using the CLA IdentiFiler™ Direct PCR Amplification Kit (ThermoFisher Scientific #A44661). PCR products were analyzed by capillary electrophoresis on an ABI 3730 Genetic Analyzer.

### 7. Karyotyping

The three iPSC lines were collected and analyzed using the KaryoStat™ assay (ThermoFisher Scientific).

### 8. Mycoplasma detection

The three iPSC lines were tested for mycoplasma using the Mycoplasma PCR Detection Kit (abmGood #G238), according to the manufacturer’s instructions.

## Discussion

Immune checkpoint inhibitor-associated myocarditis (ICIM) is a rare but potentially fatal complication of cancer immunotherapy, with reported mortality rates reaching 40–50%. Despite its clinical importance, the underlying mechanisms remain poorly understood, largely due to the lack of relevant human cellular models. In this study, we established three induced pluripotent stem cell (iPSC) lines derived from an ICIM patient, an ICI-treated patient without myocarditis, and a healthy donor. These cell lines provide a valuable platform to investigate patient-specific susceptibility to immune-related cardiotoxicity. Notably, although ICIM is a treatment-associated condition, genetic and immunological factors may contribute to individual risk, which can be preserved in iPSC-derived cell types such as cardiomyocytes.

These iPSC lines may enable the development of human-based *in vitro* models, including cardiomyocyte differentiation and co-culture systems with immune cells, to study mechanisms of immune-mediated cardiac injury. Furthermore, they may serve as a useful platform for drug screening and the evaluation of cardiotoxicity associated with cancer immunotherapies. Overall, these cell lines represent a valuable resource for advancing our understanding of ICIM and for developing safer and more effective therapeutic strategies.

## Acknowledgments

This study received funding from R01HL177581 (HZ), R01HL174432 (HZ), R01HL152055 (MM), R01HL176822 (JCW), R03HL173146 (HZ), UM1TR006031 (JCW), K08HL161405 (HZ), Stanford CVI Seed (HZ), K99CA279895 (APH), AHA Transformational Project Award (25TPA1480322, HZ), and AHA Postdoctoral Fellowship (YS, 25POST1360626).

## Data availability

Data will be made available on request.

